# Detection and quantification of introgression using Bayesian inference based on conjugate priors

**DOI:** 10.1101/2022.07.07.499145

**Authors:** Bastian Pfeifer, Durrell D. Kapan

## Abstract

Introgression (the flow of genes between species) is a major force structuring the evolution of genomes, potentially providing raw material for adaptation. Here, we present a versatile Bayesian model selection approach for the detection and quantification of introgression. The proposed *d*_*f*-*BF*_ approach builds upon the recently published distance-based *d*_*f*_ statistic. Unlike *d*_*f*_, *d*_*f*-*BF*_ takes into account the number of variant sites within a genomic region. The *d*_*f*-*BF*_ method quantifies introgression with the inferred *θ* parameter, and at the same time enables weighing the strength of evidence for introgression based on Bayes Factors. To ensure fast computation we make use of conjugate priors with no need for computational demanding MCMC iterations. We compare our method with other approaches including *d*_*f*_, *f*_*d*_, and Patterson’s *D* using a wide range of coalescent simulations. Furthermore, we showcase the applicability of the *d*_*f*-*BF*_ approach using whole genome mosquito data. Finally, we integrate the new method into the powerful genomics R-package PopGenome.

## I. Background

Our methodology builds upon the recently published *d*_*f*_ statistic, we introduced as an estimator of the proportion of introgression [1]. It is formulated as

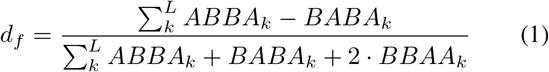

where *ABBA*_*k*_, *BABA*_*k*_, and *BBAA*_*k*_ represent SNP sharing patterns on a four-taxon tree, which we show can be expressed in terms of genetic distance:

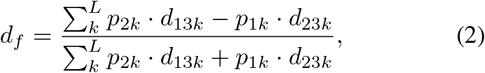

where *p*_*xk*_ refers to the mutant allele frequency in population *x* at variant site *k*. Here *d*_*xyk*_ is the average pairwise nucleotide difference between population *x* and population *y* at variant site *k. L* is the total number of bi-allelic sites in a genomic region. The first two taxa are closely related species the third taxon is a potential donor of mutant allele *B* at variable sites, and the fourth taxon refers to the outgroup as in the original work by Patterson [2]. Note, the *d*_*f*_ statistic calculates the fraction of introgression based on variant sites where the outgroup (taxon 4) is monomorphic for allele *A*.

From equation 1 it can be seen that when either *p*_2*k*_ · *d*_13*k*_ (*ABBA*_*k*_ + *BBAA*_*k*_) or *p*_1*k*_ · *d*_23*k*_ (*BABA*_*k*_ + *BBAA*_*k*_) is zero, the *d*_*f*_ statistic estimate is 1 or -1, respectively. This can generate false positives in low diversity regions, e.g in low recombining regions comprising only a few bi-allelic markers. This issue is not unique to the *d*_*f*_ statistic and applies to other *ABBA* - *BABA* methods, such as *f*_*d*_ [3], and Patterson’s *D* since they do not explicitly account for the number of bi-allelic sites which provide import evidence of introgression.

## II. New Approach

To tackle this problem, we transform the *d*_*f*_ statistic into a Bayesian model selection problem. We define two competing models of introgression.

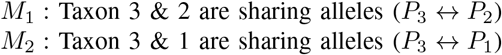

Model *M*_1_ and *M*_2_ are represented by the following binomial likelihood functions:

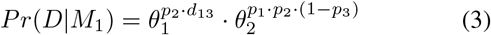

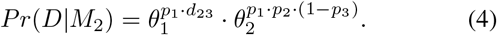

The parameter *θ*_1_ in model *M*_1_ includes information about the fraction of the data explained by the ABBA+BBAA (*p*_2_ *d*_13_) patterns. In model *M*_2_, *θ*_1_ captures the BABA+BBAA (*p*_1_ · *d*_23_) signals. The parameter *θ*_2_ includes the species tree pattern BBAA (*p*_1_ · *p*_2_ · (1− *p*_3_)), it is used as an approximate measure of the neutral (non-introgressed) signal within the data.

The proposed Bayesian model assumes that the observed data *D* can be approximately explained by the species tree pattern (BBAA) plus the corresponding introgression frequency patterns (ABBA and BABA). We use the conjugate Beta distribution as a prior

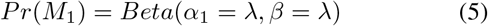

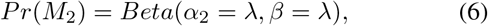

where *λ* is the average population size of *P*_1_, *P*_2_ and *P*_3_. In order to form the posterior we propose the following updating scheme of the Beta distribution per variant site *k*

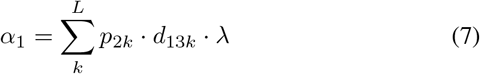

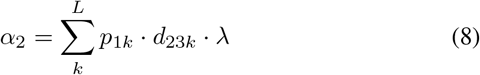

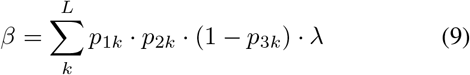

The corresponding posterior density distributions of the models *M*_1_ and *M*_2_ are

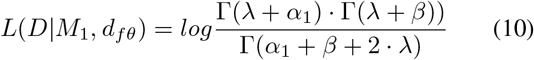

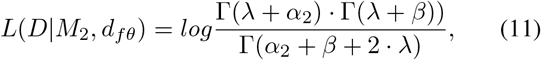

where *d*_*fθ*_ are the inferred Beta model parameter to quantify the gene-flow between *P*_3_ ↔ *P*_2_ (model *M*_1_) and *P*_3_ ↔ *P*_1_ (model *M*_2_). Finally, evidences of introgression are calculated using Bayes Factors as

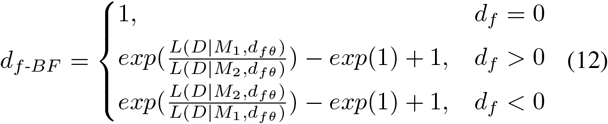

allowing researchers to judge the relative merit of the two competing introgression models. The resulting Bayes Factors are interpreted according to Jeffrey’s Table; *d*_*f*-*BF*_ = 1 (no evidence), *d*_*f*-*BF*_ = 1 − 3 (anecdotal evidence), *d*_*f*-*BF*_ = 3 − 10 (moderate evidence), *d*_*f*-*BF*_ = 10 − 30 (strong evidence), *d*_*f*-*BF*_ = 30 − 100 (very strong evidence), and *d*_*f*-*BF*_ *>*= 100 (extreme evidence).

## III. Results

To validate the *d*_*f*-*BF*_ approach we generated topologies with different levels of introgression using Hudson’s ms program [4]. The sequence alignments were produced by the seqgen program [5]. We generated 5kb sequence with split times *t*_12_ = 1 × 4*N, t*_123_ = 2 × 4*N* and *t*_123*O*_ = 3 × 4*N* generations ago. The time of gene-flow from *P*_3_ to *P*_2_ was set to *t*_*GF*_ = 0.1 × 4*N* generations ago with a fraction of introgression of *f* = 0.1. The recombination rate was set to *r* = 0.01, and a Hasegawa-Kishino-Yano substitution model was applied with a branch scaling factor of *s* = 0.01. We varied the fraction of introgression and the time of gene-flow and compared *d*_*f*-*BF*_ with Patterson’s *D* (*D*), *f*_*d*_ and *d*_*f*_. Figure 1a shows the results when varying the fraction of introgression from population *P*_3_ to *P*_2_. The *d*_*f*-*BF*_ model parameter *θ* (denoted as *d*_*fθ*_) precisely quantifies the fraction of introgression and produces almost identical results as the *d*_*f*_ statistic. The corresponding Bayes Factors for each introgression level are shown in Figure 1b. With the current setting strong evidence of introgression is reported when the fraction of introgression is greater than 0.8. We also varied the time of gene-flow. We confirm the results reported in [1], *d*_*f*_ is almost not affected by the time of gene flow, and quantifies the fraction of introgression more accurate compared to Patterson’s *D* and *f*_*d*_. We report the same properties for *d*_*fθ*_ (not shown).

**Fig. 1.**
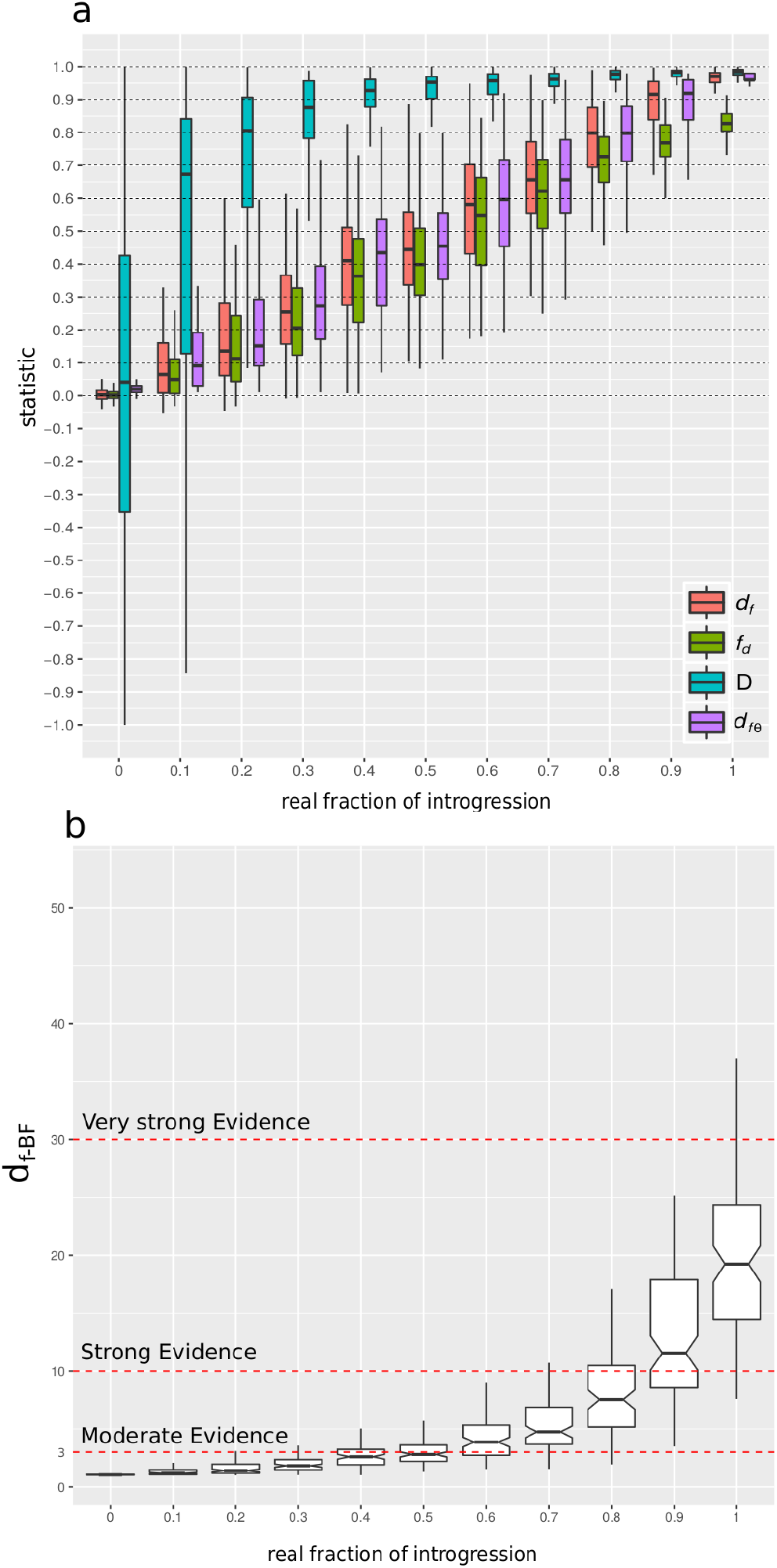
Simulation results. (a) Shown are the results of *d*_*f*_, *f*_*d*_, *D*, and *d*_*fθ*_ on simulated data with varying levels of introgression (100 iterations each). The horizontal lines refer to the real fraction of introgression. (b) The *d*_*f*-*BF*_ Bayes Factor values of the corresponding levels of introgression shown in (a).

We fully integrated the *d*_*f*-*BF*_ approach into the R-package PopGenome [6].

## Conclusion

In summary, our flexible Bayesian model selection framework quantifies introgression, is equally or more accurate than the *d*_*f*_ statistic upon which it based, and at the same time enables quantification of the strength of evidence for introgression based on Bayes Factors.

